# High-precision protein-tracking with interferometric scattering microscopy

**DOI:** 10.1101/2020.08.03.234773

**Authors:** Richard W. Taylor, Cornelia Holler, Reza Gholami Mahmoodabadi, Michelle Küppers, Houman Mirzaalian Dastjerdi, Vasily Zaburdaev, Alexandra Schambony, Vahid Sandoghdar

**Affiliations:** Max Planck Institute for the Science of Light, Erlangen, Germany; Max-Planck-Zentrum für Physik und Medizin, Erlangen, Germany; Department of Physics, Friedrich Alexander University Erlangen-Nuremberg, Erlangen, Germany; Department of Computer Science, Friedrich Alexander University Erlangen-Nuremberg, Erlangen, Germany; Department of Biology, Friedrich Alexander University Erlangen-Nuremberg, Erlangen, Germany

**Keywords:** interferometric scattering microscopy, iSCAT, iSPT, single particle tracking, live cell imaging, membrane organisation, epidermal growth factor receptor

## Abstract

The mobility of proteins and lipids within the cell, sculpted oftentimes by the organisation of the membrane, reveals a great wealth of information on the function and interaction of these molecules as well as the membrane itself. Single particle tracking has proven to be a vital tool to study the mobility of individual molecules and unravel details of their behaviour. Interferometric scattering (iSCAT) microscopy is an emerging technique well suited for visualising the diffusion of gold nanoparticle-labelled membrane proteins to a spatial and temporal resolution beyond the means of traditional fluorescent labels. We discuss the applicability of interferometric single particle tracking (iSPT) microscopy to investigate the minutia in the motion of a protein through measurements visualising the mobility of the epidermal growth factor receptor in various biological scenarios on the live cell.

## Introduction

All cells are enclosed by an outer plasma membrane and, in addition, eukaryotic cells are commonly compartmentalised by internal membranes into cell organelles, to generate specialised functional entities. The plasma membrane acts as a barrier, transport and communication platform between the cell and its environment that separates the interior of the cell from its surroundings, controls the flux of ions and nutrients and mediates sending and sensing of signals to and from the cell. These multiple functions of the membrane are enabled by its complex composition of phospholipids, sphingo- and glycolipids, cholesterol and proteins.

The plasma membrane is also a carefully regulated and highly dynamic structure which sculpts the mobility of the proteins and lipids of which it is composed (1, 2). Our view on the organisational motifs of the plasma membrane has expanded greatly over the last few decades following much investigation and advances in single-particle imaging microscopies. Present wisdom informs that the plasma membrane is organised over many length scales (1–3), with motifs including cellular domains such as cell-cell contact sites or the apical surface of epithelial cells being strictly separated from the basolateral surface of the membrane by cell-cell adhesion molecules. In addition, other motifs include cytoskeletal actin-mesh induced microscale compartmentalisation (4–6), nanoscale transient raft domains (7–9), and molecular-level crowding from protein-protein interactions (10, 11).

These organisational motifs provide a mechanism for spatial and temporal control of the lipids and proteins by hierarchically restricting their freedom of movement. Doing so allows for regions of specific composition and hence dedicated function which is required for a diverse array of physiological processes such as signal transduction, directed transport across a membrane or cell-cell-communication (1, 3). In signalling, one requires the formation of nanoscale signalling domains which comprise selected proteins. By transiently confining respective proteins, these domains are believed to regulate signalling events by changing the local concentration of proteins and as a consequence modulating efficient protein-protein interactions. In contrast, they also have the capacity to prevent signalling altogether in the absence of a stimulus through exclusionary segregation of the membrane components (1).

The ability to track the motion of proteins within the membrane on the one hand provides insights on the paths a particular protein takes during the period of observation and on the other indirectly reveals obstacles, boundaries or accelerators that affect its mobility. One thus stands to gain further insight into the functionality and structure of the membrane by observing the lateral mobility of membrane proteins as these molecules serve as ideal probes for this complex environment. Unsurprisingly, tracking protein mobility has been an important target of research over the last decades, with numerous optical techniques having been developed to meet this challenge (6). Studies at the single molecule level have shown to be extremely fruitful since they allow for direct visualisation of the behaviour of individual molecules and their molecular interactions, no matter how rare or infrequent, without fear of being lost to an ensemble observation.

Single particle tracking (SPT) of protein and lipid mobility has proven to be an exceptionally valuable and productive technique in this effort to decipher the interactions that constitute the organising principles of the membrane (12–16). SPT microscopy is typically accomplished by localising the position of the membrane molecule through attachment of an optical label, be the label a fluorescent molecule (17, 18), quantum dot (19, 20) or nanoparticle (4, 21). The purpose is then to reveal the path taken in the membrane by the molecule as detected at various temporal resolutions depending on the method of choice. While low temporal resolution can provide only partial information about the motion of the molecule without revealing the fine details of its diffusion, imaging with high frame rates effectively visualises the molecular trajectories resultant from fast and short-lived interactions within the membrane, as depicted in the illustration of **Fig. 1**.

**Fig. 1.**
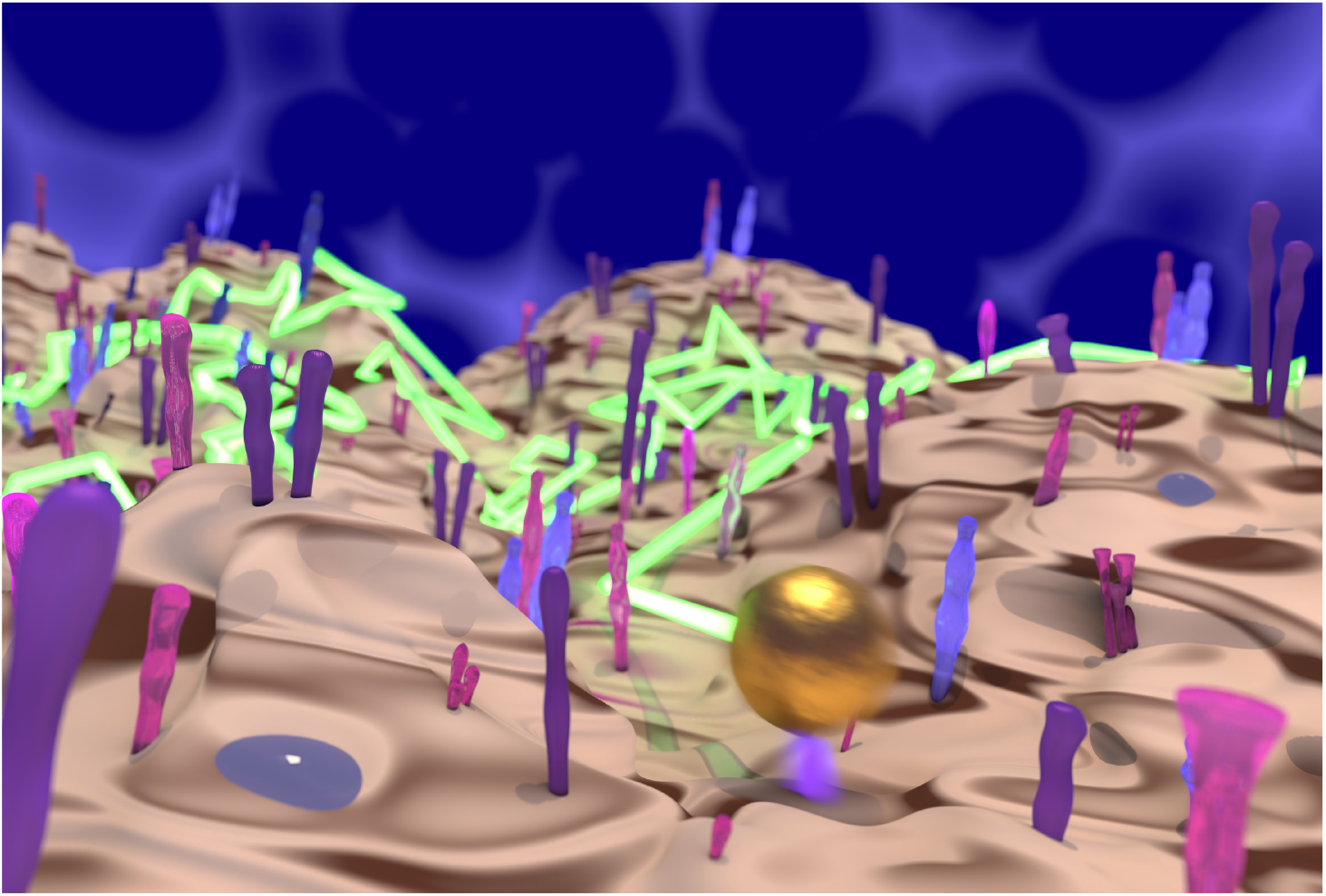
A single-molecule view of diffusion in the plasma membrane. A computer rendered illustration of a gold nanoparticle labelled-protein diffusing through the crowded terrain of the plasma membrane of the cell. Through SPT of the trajectory of the protein in all three dimensions, shown in green, one can visualise the terrain and obstacles encountered by the protein. The gold nanoparticle has the capacity to map out the local membrane topology, afforded through the nanometric three-dimensional localisation precision.

Fluorescence-based labels have been enormously popular for single-molecule imaging since they enable precise distinction between signal and background during imaging. However, fluorescence imposes several important limitations which restrict how well one can track the mobility of a membrane molecule. Firstly, the finite rate by which light is emitted from the fluorophore limits how well the position of the labelled protein can be determined. For a molecule to emit a fluorescence photon, an electron within the molecule must become promoted into a higher energy level through absorption of a photon. On relaxation of the electron, a fluorescenct photon is emitted, with the whole process taking several nanoseconds. The rate at which the label can emit photons is a critical quantity in SPT as the measured number of photons determines directly the precision to which the location of an emitter can be determined (22, 23). Since the emission rate is depending on the excitation lifetime, one therefore finds that for nanosecond lifetimes the best attainable resolutions are on the order of tens of nanometers per millisecond of recording.

In addition, fluorescent dyes are vulnerable to several processes that result in the emission of no light which is naturally detrimental for tracking microscopies. Firstly, photochemical processes can occur which alter the wavelength needed for the emitter to become excited or to fluoresce, known as *photobleaching*. Secondly, it is possible for the electron within the fluorophore to become temporarily trapped in a transient state that is dark to the illumination wavelength, a process commonly referred to as *blinking*.

The desire to gain an ever greater insight into the precise mobility of single molecules within the membrane has motivated the search for alternative optical labels that can offer improved spatial and temporal resolutions in SPT microscopy. Scattering is an attractive alternative because it does not encounter the critical limitations previously described for fluorophores. Furthermore, the permanence of scattering also enables for the scattering label to be tracked for an essentially endless period, limited only by the sample itself, and so opening the possibility to watch entire biological processes in a complete uninterrupted fashion.

The first experiments that showed the potential for scattering labels were pioneered as far back as the late 1980s (24–26). These efforts laid the foundation for the seminal investigations by Kusumi et al. whose high-speed single particle measurements revealed the existence of diffusion barriers within the plasma membrane and founded the concept of compartmentalised organisation of the membrane itself (4, 21, 27).

The challenge faced in performing SPT microscopy with nanoparticle labels is that the scattering signal is comparatively small and accompanied by a large imaging background from the remaining sample which must be tackled. Interferometric scattering (iSCAT) microscopy is an emerging technique which confronts these issues directly (28–31). With iSCAT, it recently has been shown to be possible to track the diffusion of labelled proteins on the membrane of artificial and live cells (30, 32–34) with nanometric precision in all *three-dimensions* with microsecond temporal resolution (30, 35). The recent advances of interferometric SPT (iSPT) is further spurring new efforts to understand and model the plasma membrane (36).

In this work, we show how iSCAT microscopy is highly suited for investigating the mobility of a transmembrane signalling protein to fine detail within the live cell membrane. We chose the epidermal growth factor receptor (EGFR) as a candidate since EGFR is a prototypical receptor tyrosine kinase belonging to the ErbB family and is a key regulator of cell proliferation, growth, survival and differentiation in mammalian cells (37). EGFR is an 1186 amino acid single-pass transmembrane glycoprotein that binds and is activated by ligands of the Epidermal Growth Factor (EGF)-family with 11 known members in humans (38).

In its inactive state, EGFR is mostly residing and travelling within the plasma membrane either as a monomer or as preformed but inactive dimers, with the ability to continuously switch between these two states (39). To be primed for signalling, however, EGFR must be in the dimerised state in order to become active after EGF ligand binding. Signalling is initiated by autophosphorylation of the intracellular part of the receptor, which constitutes binding interfaces for signalling proteins from which the signal is further transduced along the signalling cascade. Once activated, EGFR becomes rapidly endocytosed and continues signalling in endocytic compartments until the receptors either are returning to the inactive state and recycled back to the membrane or become degraded via the lysosomal pathway (38, 40–45).

Since EGFR signalling strongly depends on receptor mobility as outlined above and aberrant signalling of EGFR is associated with various cancers by promoting oncogenic signalling, this receptor has been widely investigated as a model system to explore how the nanoscale organisation of the plasma membrane affects signalling function (37, 46). For this reason, EGFR has also served as a model membrane protein for numerous SPT microscopies (21, 39, 47–50).

Here, we show how the high spatial and temporal resolution of iSPT can be used to uncover fine details in the trajectories of an EGFR in scenarios such as diffusion in the membrane as well as applicability in investigating processes such as endocytosis and trafficking. We discuss several methods for statistical quantification of single particle trajectories that take advantage of the high temporal resolution to reveal detailed information about protein mobility in the membrane such as nanoscale sub-diffusion and confinement events. We first begin by introducing the principle of iSPT microscopy and in particular how three-dimensional (3D) nanometric particle localisation is accomplished.

## iSPT on the live cell

### A. SPT microscopy: interferometric scattering vs fluorescence

In fluorescence-based SPT, one follows the molecule of interest by detecting the light emitted by an attached fluorescent label. Since the light emitted by the label is shifted to a longer wavelength than that used for excitation, one can exclusively detect the labelled protein through the use of convenient wavelength-selective filters. This detection strategy has proven enormously successful, with single-molecule sensitivity now being something widely accessible to many laboratories.

However, the aforementioned inconsistent and low number of photons emitted through fluorescence curtails the resolution and fidelity by which one may identify the position of the target molecule. This problem is further compounded with the presence of detection noise which is often of a similar intensity to the weak probe signal.

Light scattering is an alternative optical process to fluorescence which overcomes many of these difficulties. In light scattering, the incoming light briefly couples to the electrons in the material before being swiftly re-radiated away. For this reason, whilst not all materials can fluoresce, all are able to scatter light. Moreover, light scattering is not vulnerable to the processes of blinking and bleaching as encountered with fluorescence.

One advantage of light scattering is that the interaction between the photon and the electron is near-instantaneous. For a metal nanoparticle, such as gold, this interaction can be as fast as 10 fs and hence one can generate a million scattered photons in the same time it would take to generate a single fluorescence photon.

The process of light scattering also has the crucial property of being coherent, a feature not shared by fluorescence. In being coherent, the light scattered by the particle retains the same temporal signature as the illuminating light, meaning they share the same phase difference throughout time. Hence, two light beams can be meaningfully added together in an optical process known as interference. The advantage in interfering two optical beams together is that if one of the beams is weak in amplitude - for example as occurs with nanoscale scattering, then one can boost the effective amplitude of this weak signal by mixing with a stronger coherent companion beam. Oftentimes one can boost the weak scattered signal far above the level of normal detection noise, thus offering exquisite detection sensitivities. Microscopies that harness this detection principle are described under the umbrella term ‘interferometric microscopies’, and in particular iSCAT is a technique that has been developed to optimise the sensitivity for the detection of nanoparticulate matter.

We can express the interferometric detection of scattered light via the following equation:

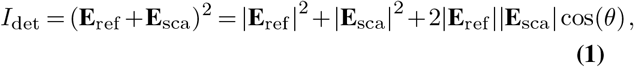

where *I*_det_, is the light incident upon our detector, **E**_sca_ the electric field of the scattered light of the sample and **E**_ref_ is the companion beam with which we interfere the scattered light. In practice this beam, referred to as the ‘reference’ beam, can be obtained from a reflection of the illuminating beam in the optical path, which we discuss further in the following section. The key component of Eq. 1 is the cross-term where one sees the multiplication of the scattered and reference fields to create a product with an overall larger intensity.

The phase term cos(*θ*) determines the contrast of the interference term with respect to the bright signal of the reference intensity and the exact phase difference is determined by many contributions including the material of the scatter, the imaging arrangement and sample geometry (35).

*niSCAT microscopy on the live cell A popular modality for iSCAT microscopy is the wide-field reflection arrangement shown in **Fig. 2(A)**, where the sample is illuminated by a wide beam, and the reference beam (labelled **E**_ref_) with which the sample-scattered light (**E**_sca_) interferes originates from the portion of this incident illumination which back-reflects off the sample-bearing glass coverslip.

**Fig. 2.**
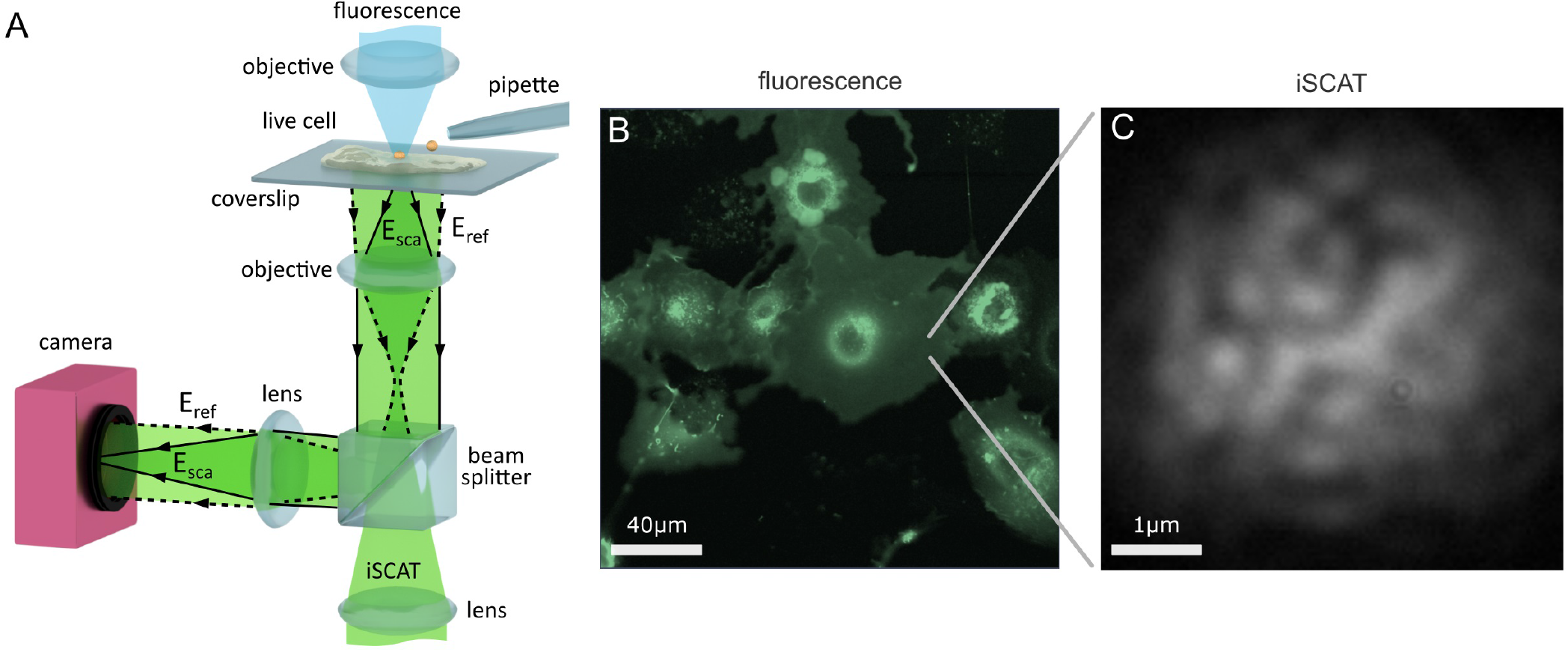
Imaging the live cell with iSCAT microscopy. **(A)** Schematic of a wide-field reflection iSCAT microscope which incorporates a micropipette and a confocal fluorescence imaging channel. **(B)** A complementary macroscopic confocal fluorescence image of COS-7 cells transfected to over-express the EGFR-EGFP protein. **(C)** A raw widefield iSCAT image of the COS-7 membrane from the region highlighted in **(B)**.

A key advantage of iSCAT microscopy is that the high sensitivity of detection permits imaging with excellent signal-to-noise levels even at extremely high speeds, with framerates in the range of 100-1,000,000 frames per second (fps) reported in the investigation of membrane diffusion (30, 32–34, 51–53). For imaging protein mobility on live cells, high framerates are particularly important in order to observe transient nanoscale molecular interactions which occur at swift milli-to microsecond time scales that may otherwise be not observable at lower framerates (54). Illumination intensities required for these high framerates are typically in the order of I = 1*−*10 kWcm^*−*2^, which are compatible with live cell imaging (55).

In this work, we perform our investigations upon live HeLa and COS-7 cells. HeLa cells are an adherent, epithelial cell line derived from human cervix tissue and COS-7 cells are an adherent but fibroblast-like cell line that originate from the kidney tissue of the African Green Monkey. On HeLa, we track the endogenous EGFR whereas on COS-7 we track exogenous EGFR. In the latter case, this is achieved by transfecting the COS-7 cells with a plasmid encoding EGFR which increases the level of EGFR expressed within the cell. We link the EGFR plasmid with the enhanced green fluorescent protein (EGFP) to yield the fusion protein construct EGFR-EGFP which then allows us to visualise successful transfection through the fluorescence signal of EGFP. An asset of iSCAT microscopy is that it is fully compatible with fluorescence imaging (56), and so one can incorporate conventional fluorescence microscopies and labelling strategies to assist with investigation on live cells.

We introduce a scanning confocal fluorescence channel from above the sample, shown in **Fig. 2(A)**, to provide a macroscopic overview image of the live cell sample, as shown in **Fig. 2(B)**. iSCAT microscopy may also be performed in parallel to confocal fluorescence imaging, and **Fig. 2(C)** presents an image of a portion of the lower cell membrane viewed with iSCAT microscopy. The nature of iSCAT microscopy, particularly in this back-reflection scheme, produces interferometric images of the cell membrane that show a characteristic speckle pattern as shown in **Fig. 2(C)**.

#### On labelling in iSPT

The universality by which all materials scatter light results in their ability to be detected in iSCAT microscopy, provided appropriate care is taken to mitigate optical noise associated with the detection of light. In particular, it indeed has been shown to be possible with iSCAT microscopy to detect individual proteins directly without the need for any label when observed in isolation upon glass coverslips (57–59).

In tracking lipids and proteins when incorporated into a membrane, it is necessary to invoke use of a label to identify the molecule of interest and to render it distinct. Just as one can use a fluorescent label, one can equally use a scattering label. Gold nanoparticles (GNPs) are widely used scattering probes owing to their excellent scattering efficiency and bio-compatibility. As with fluorescence labelling, use of a colloidal label can potentially introduce artefacts that affect the diffusion under study, and these issues are addressed later in the discussion section.

#### 3D localisation with iSPT microscopy

In iSPT, one images the light scattered by the GNP-labelled protein. However, since the remaining cell and its constituents also scatter light, they also produce a signal which we refer to as the imaging background. To localise the GNP-labelled protein in any given frame, it is first necessary to isolate this probe signal out of the imaging background.

Removal of the imaging background of the cell is, in general, challenging (29). This is because firstly the scattered light has the same wavelength as the illumination and sample background and so simple wavelength-selective filtering is not possible. Secondly, the cell background is dynamic and fluctuating in nature, and consists of features with fine details of a similar size and shape to the probe signal. This dynamism stems from the fluctuating composition and morphology of the membrane and the cell interior, which is then observed as a random speckle-like pattern with temporal fluctuations in intensity, as illustrated in **Fig. 2(C)**.

Several strategies have recently been reported which address this problem of removing the interferometric imaging background of the cell (34, 60). One strategy exploits the difference between the highly symmetric circular pattern of the imaged gold probe and the ‘random’ pattern of the cell background speckle (30). **Figure 3(A)** presents an image of the plasma membrane of a COS-7 cell, where a single GNP has become bound to the membrane. As the nanoparticle is smaller than the optical diffraction limit, in imaging the GNP, especially interferometrically, one sees not an image of the particle but instead a series of rings of alternating contrast referred to as the interferometric point spread function (iPSF) (35). By using a circularly-symmetric feature extraction algorithm, the probe iPSF can be read from one frame of the video (30, 35). The imaging background, however, should not necessarily be viewed as a nuisance, rather it can also provide useful information. For example, the interferometric image of the cell can indicate sample creep and drift which are important parameters to quantify and correct for in wanting to reliably interpret single particle trajectories (30). Once the probe iPSF has been isolated from the background, one can then determine to nanometric precision the 3D position of the probe. The lateral (*x-y*) position is found by determining the centre of the symmetric iPSF, which can be accomplished through a variety of means such as by finding the centre of symmetry (61), or through fitting of a two-dimensional Gaussian intensity function or an interferometric diffraction-based model which describes the iPSF exactly (35).

**Fig. 3.**
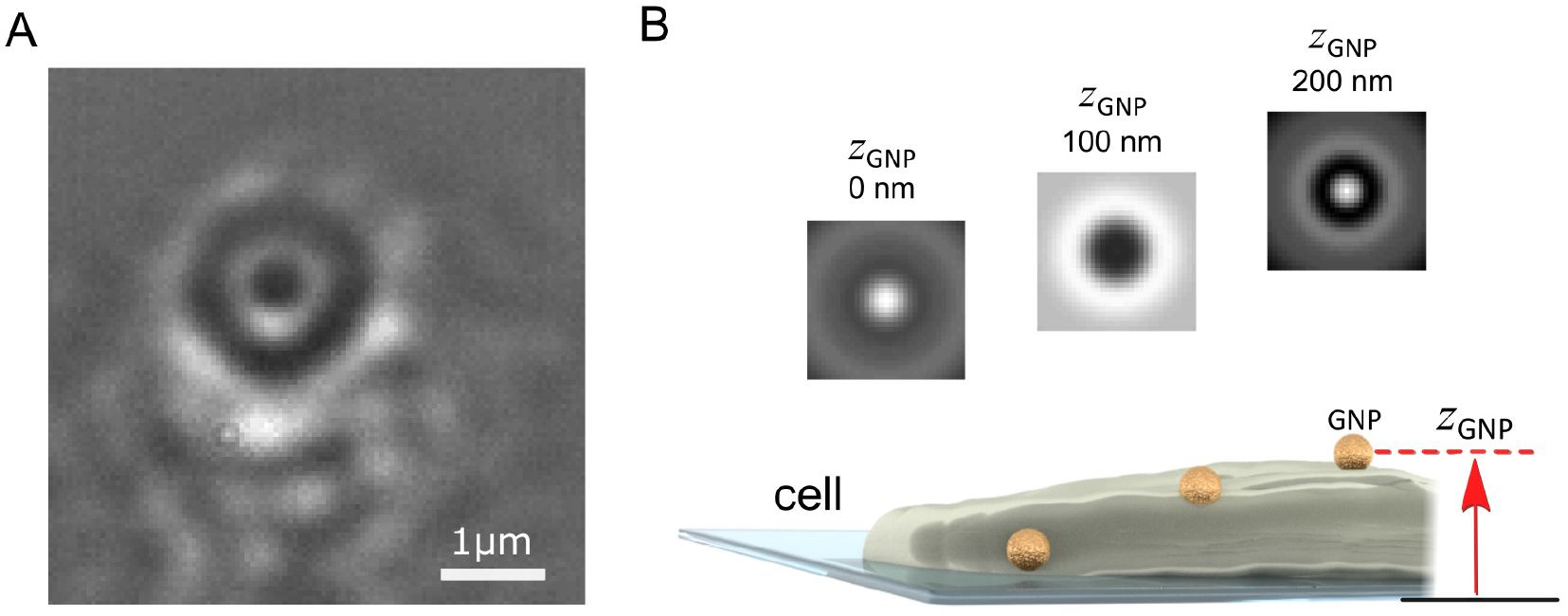
Imaging and localising a GNP probe in 3D with iSPT. **(A)** Background-corrected iSCAT image of the COS-7 cell membrane with a bound GNP probe (diameter 48 nm). The GNP appears as a series of black and white rings from the iPSF. The distinctive ring pattern depends on the height of the GNP probe above the coverslip and also its position relative to the objective focus. **(B)** Schematic illustrating how the height of the GNP probe enables determination of its axial position: as the GNP diffuses on the membrane, a change in the height of the GNP above the coverslip produces a distinct iPSF which can be used to determine the height of the GNP *z*_GNP_.

To determine the height of the GNP above the coverslip (*z*_GNP_), one exploits the information encoded within the alternating contrast of the iPSF ring pattern, which is a distinctive feature of iSPT (35). **Figure 3(B)** illustrates several iPSFs for a GNP that is diffusing over a region of the cell membrane which raises its height. The distinctiveness of the iPSF and the good sensitivity by which it can be measured enables nanometrically-precise lateral and axial localisation. One can determine the relative change in height by direct calibration of the central contrast of the iPSF (62), or by machine-learning assisted clustering of the iPSF (30) or by direct fitting of the iPSFs to a model specifically describing iPSF formation (35).

#### Watching *in situ* landing of a GNP probe on the membrane with iSPT

To provide an illustration of 3D nanometric iSPT of a GNP probe, we present **Fig. 4**, wherein we track the landing of a GNP probe upon the cell membrane. To observe the labelling step is an important ability in interferometric particle tracking as one is able to ensure that indeed it is the GNP probe which is being observed but also one is able to know when the probe-protein interaction began which is especially important for investigating time-dependent processes.

**Fig. 4.**
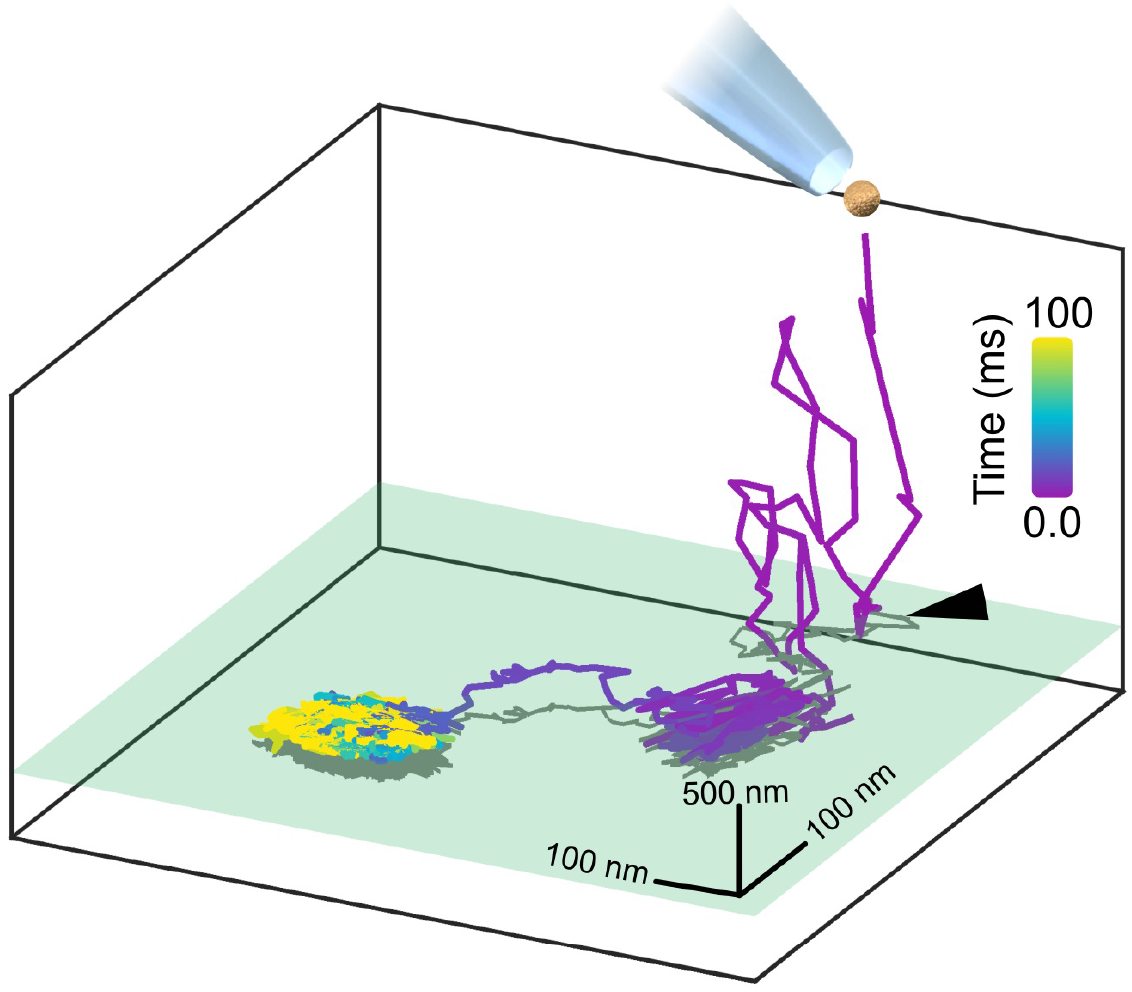
Watching *in situ* the moment a GNP lands upon the plasma membrane. A GNP, once pipetted towards the COS-7 cell, is tracked axially over a range of 2 *μ*m whilst it diffuses in the culture medium before coming to land upon the cell membrane, with the initial point of contact with the membrane marked with a black arrow. The axial plane of the cell membrane is represented in green and a two-dimensional projection of the trajectory is plotted in grey. The whole landing event lasted approximately 100 ms and was recorded at 30,000 fps. The GNP used here was positively-charged so as to immediately and readily bind non-specifically to the negatively-charged cell membrane for the purpose of illustration.

**Figure 4** demonstrates the *in situ* binding of a pipetted GNP probe to the plasma membrane, visualised through the 3D trajectory. Here in this example, for illustration we use a positively-charged GNP which readily binds to the negatively-charged plasma membrane. It should be noted that in this case, binding to the plasma membrane is purely electrostatic and not representing any particular lipid or protein motion in the membrane. Here, the GNP is tracked axially over a range of two microns where it is then seen to initially interact with the membrane, break loose before then ultimately becoming permanently attached.

#### Describing the erratic mobility of proteins

If one watches the path taken by an individual protein as it navigates through the environment of the cell, one sees the protein takes steps that are seemingly random in length and direction. This behaviour is driven by ever-present random fluctuations in the environment of the protein and is referred to as diffusion. When the mobility of a particle is driven by thermal fluctuations and otherwise unencumbered, we refer to the mobility of the particle as being Brownian or ‘normal’. When normal diffusion is enhanced or restricted by an external influence, for example, by a variety of geometric and steric interactions, then the diffusion becomes anomalous.

Anomalous diffusion encapsulates two regimes of mobility that are in many regards polar opposites. Anomalous diffusion, wherein the mobility is enhanced by the presence of phases of persistent motion, for example by the application of an external driving force, or complex velocity flow, is referred to as super-diffusion. Conversely, anomalous diffusion that is generally hindered or restricted by the presence of, for example, traps, obstacles and barriers, is especially referred to as sub-diffusion (63–66). In the plasma membrane, sub-diffusion results from interactions with the organisational elements of the membrane, in the first place lipids and other proteins in or attached to the membrane, which constitute domains, barriers or corrals (2, 10, 11). Consequently, investigations into and determination of sub-diffusion remains an active area of research to understand membrane activity, structure and function (67, 68).

##### Mean square displacement analysis of membrane mobility

The general diffusive nature of membrane protein mobility implies that it is impossible to predict how the protein will proceed from one step to the next as the process is essentially random. Nonetheless, various statistical models have been proposed to interpret these random walks in order to describe any characteristic properties they may have (69, 70). One immediate characterisation is, for example, whether the walk exhibits normal or anomalous behaviour, which may be an important indicator of function in the case of a transmembrane protein.

One of the most widespread means of analysis is that of the Mean Square Displacement (MSD) where one examines the time-dependent character of the diffusive walk to quantify whether the walk is normal or anomalous, and if it is anomalous, to which degree. We can express the MSD as:

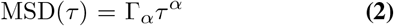

where *τ* represents an interval of time within the trajectory, Γ_*α*_ is a coefficient of proportionality and *α* is the temporal exponent (time-dependence) that we seek in order to quantify the nature of the diffusive walk. A protein undergoing unencumbered Brownian diffusion is characterised by a MSD which is linear in time, i.e. *α* = 1, which is so-called ‘normal’ diffusion and Γ_*α*=1_ = *D* is just the diffusion constant. For a diffusing protein that is not truly free, i.e. being subject to additional influences beyond thermal fluctuations, then the diffusion can be anomalous and *α* ≠ 1. An *α* > 1 describes super-diffusion, and occurs, for example, in instances where the protein is being specifically guided along a certain path such as which occurs in intracellular transport. An *α* < 1 denotes the hindered mobility of sub-diffusion, and for diffusion measured in the plasma membrane a typical value of *α* = 0.7 is characteristic by conventional ensemble MSD analysis (67). With careful analysis of the MSD and temporal exponent, one can hence characterise the diffusive character of the membrane protein within the local membrane environment.

To compute the MSD, one calculates how the average distance walked squared for a given interval of time depends on how long the time interval is. To obtain a reliable measure of the time-dependance of the walk one requires a sufficient number of recordings to give an average that is statistically robust (71–73). One method is to build an average out of many similar trajectories from separate measurements. Alternatively one may take a single sufficiently long trajectory and subdivide it into shorter pieces which are then averaged together. The former approach is known as a *particle* ensemble and the latter is referred to as a *temporal* ensemble. A diffusive system in which these two approaches are equivalent is said to be *ergodic*. The plasma membrane of the cell is an example of a system that can exhibit weak ergodicity breaking which is a behaviour chiefly originating out of inhomogenous confinement and stalling of the diffusing protein in the membrane (69). As a consequence, these two approaches can not generally be assumed to be equivalent.

In our work presented here, we compute the temporal average. Furthermore, since one can accumulate a sufficient number of frames from a very short window in time, typically representing just a small fragment of the complete trajectory, one can roll the MSD analysis incrementally across the whole trajectory and thus build up a measure of how the temporal exponent is changing throughout the trajectory. Doing so allows one to visualise how the mobility is evolving in time as the diffusing protein encounters new and differing obstacles and environments (30). The rolling window time-averaged MSD is given by:

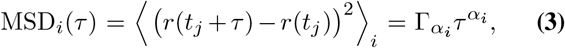

where *i* denotes the index of the temporal window wherein the MSD is to be computed, *r*(*t_j_*) is the *x*-*y* position of the protein at a time *t_j_*, where *t_j_* represents points in time belonging within the window and ⟨…⟩ denotes the calculation of an average. The temporal exponent which characterises the time-dependence of the MSD within the given window *i* is denoted by *α_i_*.

The microsecond imaging resolution of iSPT allows one to quantify the time-dependence of the protein mobility to millisecond resolution when employing the rolling temporal average. One, however, must also be careful when analysing trajectories recorded at high-frame rates. Firstly, in general, one must always account for the localisation precision for each specific trajectory point (*σ_xy_*) in the computation of the MSD as an omission of this uncertainty would lead to under-estimation of the respective temporal exponent (74). When imaging at high framerates in particular, it is possible that the step taken by the protein between two pairs of frames is comparable to, if not smaller than, the certainty with which the start and end positions are able to be known. In such cases, when performing the analysis one must be careful to avoid use of these frame pairs to avoid erroneous calculation of *α_i_*.

##### Directional correlation analysis

A complementary means to quantitatively identify the presence of obstacles and barriers to diffusion or directed transport is to look for correlation in the direction in which any two steps are taken. In the case of free diffusion, one would expect no correlation in the walked trajectory as the process is essentially random with no memory. In the presence of obstacles, however, one would expect the occurrence of recoil or ‘knock-back’ upon collision with said obstacles. In the case of directional transport, one would expect a prevalence of a certain direction, i.e. a positive correlation. Directional correlation analysis seeks to quantify the occurrence of such collisions by considering the angular correlation between two steps, and seeing if this is a meaningfully repeated occurrence.

The directional correlation is calculated by first considering two steps taken by the diffusing probe. The first step, 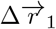, is the vectorial displacement between two frames separated in time by 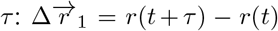, where here we take *τ* = 5 frames. The second step begins where step 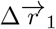 ended, and also occurs over an equal time interval, i.e. 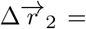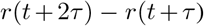. The cosine of the angle between these two steps is then computed, thus giving an expression for their angular correlation. As with the MSD, this calculation is repeated for all pairs of steps that occur within a smaller window (*i*) of the whole trajectory so that changes in the directional correlation can be identified over the course of the trajectory. Formally, we calculate the directional correlation *C_i_* for the vectorial steps 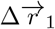 and 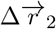 via the following (75, 76):

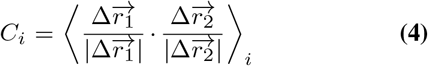

where |…| denotes the magntiude of the vectorial step and as before, the average is computed for all pairs in the *i^th^* window. A truly random process possesses no overall angular correlation between any pairs of steps, and so *C* = 0. A positive directional correlation would imply the diffusing probe exhibits some persistent motion and a negative correlation suggests persistent knock-back or deflection.

## Tracking EGFR on the live cell with iSPT

### Labelling the receptor

In our investigation to label the EGFR, we functionalise GNPs of 48 nm diameter with EGF to produce an EGF-GNP probe that specifically binds to the EGFR. To do so, we use streptavidin-coated GNPs and EGF probes with attached biotin molecules. The strong chemical bond between streptavidin and biotin serves as the linker between the GNP and the EGF ligand. A micropipette is then used to deliver small quantities of the EGF-GNPs to the immediate vicinity of the cell under investigation. This introduces the distinct advantage of setting a clear start point in which the probe is known to begin to interact with the membrane, which is important for biological processes stimulated by probe binding. The permanence of the scattering signal from the GNP probe along with the ability of iSPT to track nanoparticles in 3D over an extended axial range permits observation of the labelling process prior to immediate tracking of the mobility of the labelled protein. Once the functionalised GNP has bound the receptor we refer to this complex as ‘EGFR-GNP’.

### B. Anomalous mobility on the plasma membrane

In **Fig. 5** we present an example of a typical trajectory following the labelling of the EGFR with an EGF-GNP probe. This trajectory was recorded on the lamellapodium of a COS-7 cell at a framerate of 30,000 frames per second for a duration of 8.3 s (250,000 trajectory points). The *x-y* projection of the trajectory is plotted in **Fig. 5(A)**. The axial information of the trajectory reveals diffusion on a locally-flat but slowly curving surface expected from the lamellapodium, depicted in **Fig. 5(B)**.

**Fig. 5.**
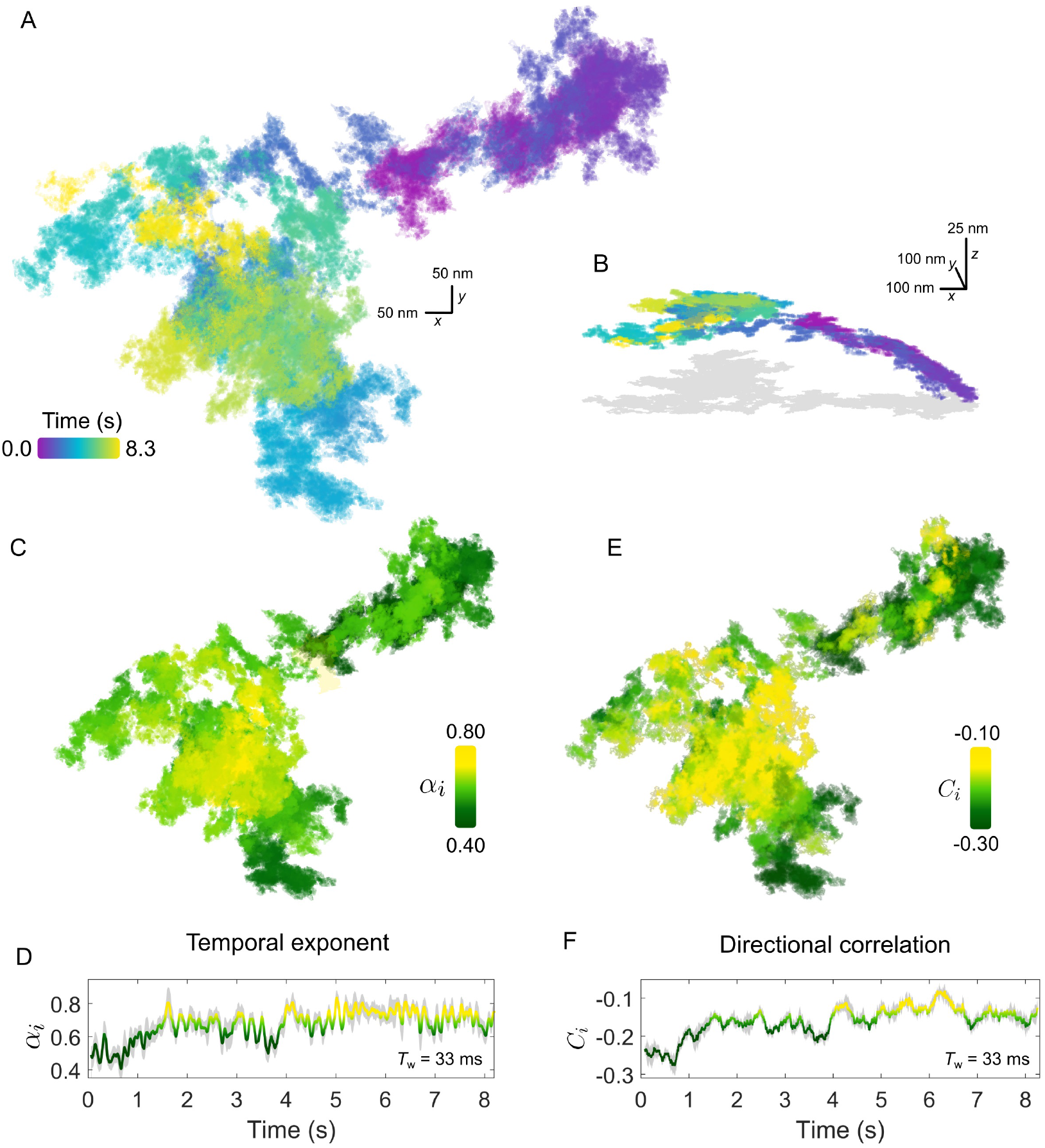
Anomalous EGFR diffusion in the plasma membrane revealed by high-speed iSPT. **(A)** A typical EGFR-GNP trajectory recorded with 30,000 frames per second for a duration of 8.3 s on a COS-7, here shown as *x-y* plane. **(B)** The trajectory from **(A)** viewed in 3D, revealing global curvature. The 2D projection is depicted in grey. **(C)** The rolling temporal exponent of diffusion *α_i_* for the trajectory for an incrementally sliding window of 1,000 frames (*T*_w_ = 33 ms), represented by the colour scale, mapped onto the *x-y* trajectory. **(D)** The temporal exponent plotted as a function of time. **(E)** Similar to **(C)**, the rolling directional correlation *C_i_* is mapped onto the *x-y* trajectory and also expressed as a function of time **(F)**. The grey shadow in **(D)** and **(F)** show the bounds of uncertainty in calculation of each respective quantity.

The extended period of observation shows the frequent occurrence of the probe revisiting the same regions of the membrane as well as navigating through distinct patches with differing sparsity of trajectory points. One must express care in interpreting the trajectories by visual inspection, and thus we defer to quantification by rolling MSD and directional correlation analysis. Firstly, by setting the rolling window to a length of 1,000 frames, corresponding to a window with temporal duration *T*_w_ = 33 ms, we are able to explore the ‘microscopic’ changes in the diffusional character of the trajectory. The temporal exponent of diffusion is plotted in **Fig. 5(C) and (D)**. Generally, the trajectory has a global mean value of *α* = 0.7 *±* 0.1 confirming indeed the diffusion is anomalous. Inspection of the rolling value reveals a distribution in the value of *α_i_*, which appears to be region-specific, with some portions of the walk encountering strong confinement with *α_i_ ≈* 0.5. In comparing the directional correlation, **Fig. 5**(**E**) and **(F)**, to their respective temporal correlation partner, we find strong agreement between both methods of analysis with the same temporal fluctuations and spatial distribution evident in both. The mean directional correlation value for the trajectory, *C* = *−*0.2 *±* 0.1 implies deflection and knock-back of varying strength. In tandem, both *α* and *C* suggest that the sub-diffusion occurring in the trajectory results from the obstructed random walk of the EGFR-GNP, and that this interaction is locally heterogeneous.

The origin for the heterogeneous environment encountered by the EGFR protein - which manifests in the anomalous trajectory of the protein - is multifold. Whilst it is beyond the scope of this work to give a comprehensive account of all the known means of interaction, it is nonetheless interesting to consider some of the core principles of membrane organisation that readily affect EGFR. To that end we might begin with the role played by the substructure and compartmentalisation of the plasma membrane itself. Kusumi and co-workers first proposed that the membrane is partitioned into compartments, sometimes doubly so, by the underlying actin cytoskeleton which thus acts as a diffusive barrier to membrane proteins, with compartment side lengths in the range *L* = 40-800 nm depending on the cell type (4, 5, 21, 27, 77). These compartments, which frustrate free diffusion, appear to render useful biological function. For example, monomeric proteins tend to be able to pass the compartment boundaries more easily via a process known as ‘hopping’ or ‘hop-diffusion’, whereas dimeric or oligomeric forms of the protein tend to remain confined (77). As a result, such compartments may enrich or exclude multimeric proteins and thus constitute functional platforms in e.g. signalling or adhesion.

It is interesting to reflect upon the fact that our understanding of the important role membrane compartmentalisation plays upon diffusion was borne out of the early pioneering work of Kusumi and co-workers more than two decades ago. By using one of the earliest realisations of iSPT microscopy which granted them sufficient temporal resolution, membrane compartmentalisation and ‘hop-diffusion’ could be inferred through careful analysis of the trajectories of membrane proteins. With the recent progress in improved detection sensitivity, spatio-temporal resolution and precision from the ever-expanding iSCAT and iSPT microscopy community (30, 33, 57, 59), it is an exciting prospect to revisit and explore the role membrane compartmentalisation plays on protein mobility with renewed experimental and theoretical vigour (36).

We next consider an additional example of membrane diffusion similar to **Fig. 5**, shown in **Fig. 6**, where the lateral trajectory (**Fig. 6(A)**) shows again a heterogeneity and sparsity in the walked path as well as clear re-visiting of domains as opposed to the continued visiting of new membrane areas. Performing the MSD **(Fig. 6(B,C)** and directional correlation analysis (**Fig. 6(D,E)**) upon this trajectory for again a window of 1,000 frames (*T*_w_ = 33 ms) reveals interesting features. Firstly, we observe again strong heterogeneity in *α* and *C* throughout the trajectory that is region-specific but not necessarily always the same through the course of time of the measurement. In particular, the temporal exponent shows a strong variance in value but generally is centred around *α* = 0.7. Interestingly, the directional correlation shows general agreement in trend with the temporal exponent, and is also suggesting ‘knock-back’. But unlike the example of **Fig. 5**, there is weaker correspondence between the sharp changes in the degree of sub-diffusion and the step correlations. Since these two forms of analysis probe different aspects of the EGFR-GNP trajectory this is entirely plausible. One possible interpretation is that the labelled EGFR-protein is slowed down within a ‘sticky’ sub-domain because then the distance travelled in time, quantified by *α_i_*, would show sub-diffusion, whilst if no immediate barrier or boundary is encountered no correlation in steps would be evident. Here we define sticky to mean orders of magnitude longer residency than typically observed from the surrounding membrane, here longer than 1 ms.

**Fig. 6.**
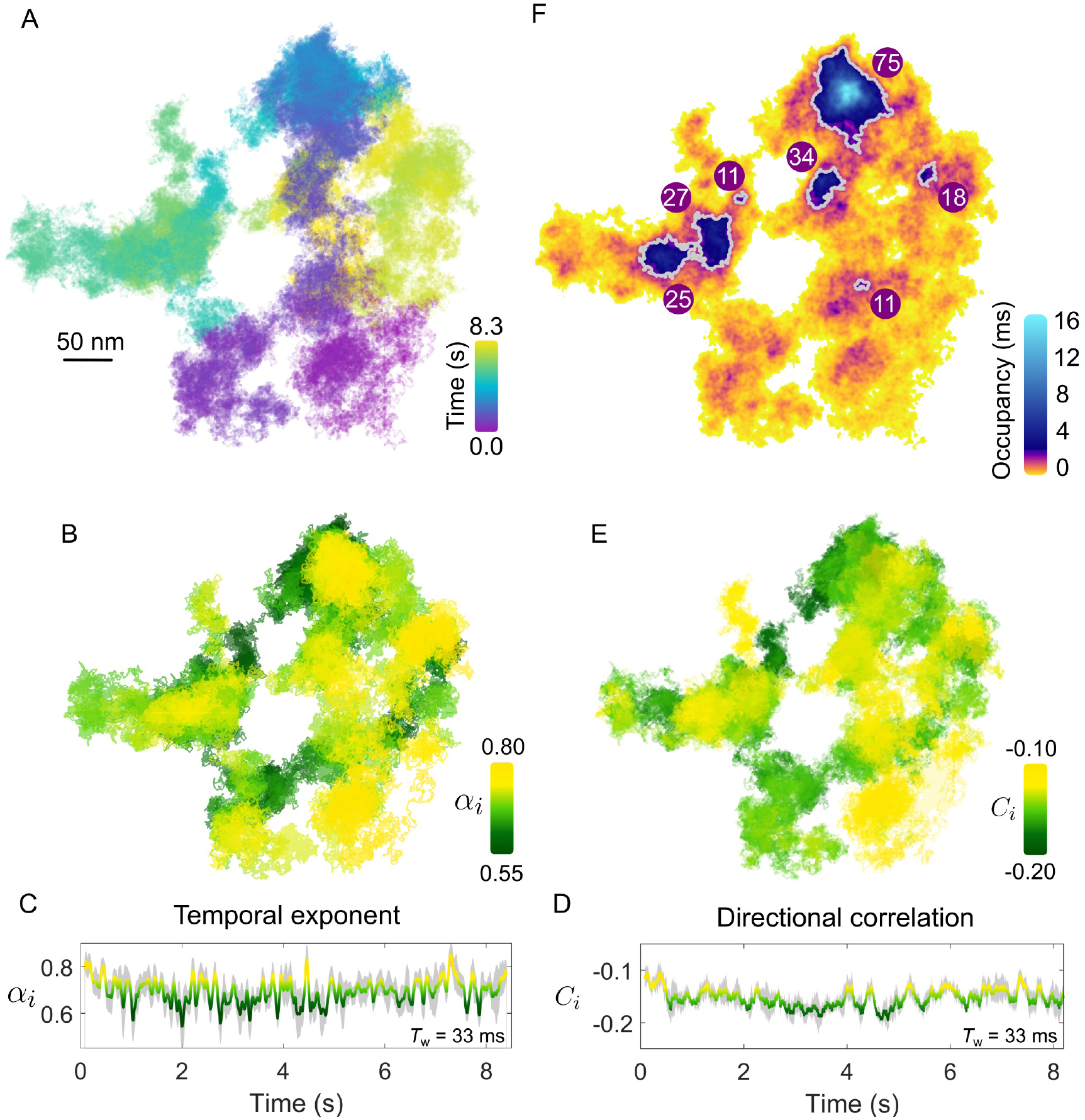
Confining patches within an anomalous walk. **(A)** The EGFR-GNP trajectory projected in x-y plane recorded at 30,000 fps over 8.3 s on a COS-7 cell. (**B**) The temporal exponent of diffusion is plotted for the trajectory as the colour-scale over the trajectory showing the spatial distribution. **(C)** The temporal exponent of diffusion plotted as a function of time. **(D)** The directional correlation is also plotted as a function of time. (**E**) Directional correlation plotted over the corresponding position in the trajectory. and time . **(F)** The density of trajectory points across the space of the trajectory is plotted as an ATOM map where colour denotes the occupancy of each 2D areal bin of size 4*×*4 nm^2^. Patches of extra-ordinary occupancy are outlined in grey as a guide to the eye. The numbers plotted represent the equivalent circular diameter in nm of the outlined patches.

Furthermore, visual inspection also reveals the presence of patch domains consisting of a higher density of trajectory points. The presence of nanodomains on the membrane, where one believes that diffusion of a transmembrane protein becomes transiently arrested, is of great interest. The high temporal resolution of iSPT permits us to visualise the presence of domains where the EGFR-GNP becomes confined by simply dividing the *x-y* trajectory into 2D bins and counting the occupancy of each bin. Such a plot we refer to as the accumulated total occupancy map (ATOM) (30). Confining domains would consist of a higher population of trajectory points than non-confining, sparse regions. **Figure 6(F)** presents the ATOM plot for the trajectory for an areal bin size of 4*×*4 nm^2^. Here we use a highly non-linear colour scheme to denote the occupancy to elucidate the wide range in residence times. We calculate the residence time by multiplying the total number of trajectory points per bin by the exposure time of one frame. The ATOM plot of **Fig. 6(F)** displays several clear patches of B Anomalous mobility on the plasma membrane extended residency with a mean equivalent circular diameter in nanometres illustrated on each prominent patch outlined in grey. Such regions of extended residency might represent favorable biochemical, structural or morphological features of the cell membrane, although lacking further information one can only speculate on the nature of such features.

One advantage of iSPT for investigating membrane processes is the ability to observe the EGFR-GNP for an extended, in principle indefinite, period owing to the absence of photo-degradation of the GNP probe. Coupled with the high temporal resolution and the sliding window analysis one can therefore monitor cellular processes in which the EGFR-GNP is involved, as reflected through marked changes in its diffusional behavior.

**Figure 7** presents such an example for diffusion occurring on the plasma membrane. Inspecting the lateral trajectory **Fig. 7(A)** reveals an apparent change in the diffusion which in time becomes progressively more spatially confined, with eventually a concerted directional motion replacing the random-like exploration of the membrane. Examining the 3D projection of the trajectory (shown in the inset of **Fig. 7(A)**), one sees the latter directed phase of the diffusion carries the EGFR-GNP down and away from the region of initial confinement. These distinct changes also manifest clearly in the kinetic parameters of the trajectory. For example, the temporal exponent (**Fig. 7(B,C)**) confirms a stark transition from a nearly-free like diffusion (*α_i_* = 0.9) in the first several seconds of observation that quickly transitions into a strongly confined mode (*α_i_* = 0.4) that is maintained until the end of observation. Our sliding window analysis explores the diffusive nature to time lags bounded within the window length, here *T*_w_ = 167 ms, and thus we discover that within this phase the EGFR-GNP is strongly confined. These observations are similarly reflected in the directional correlation which displays persistent recoil from a confining obstacle and identifies three stages of the trajectories evolution, marked (i)-(iii).

**Fig. 7.**
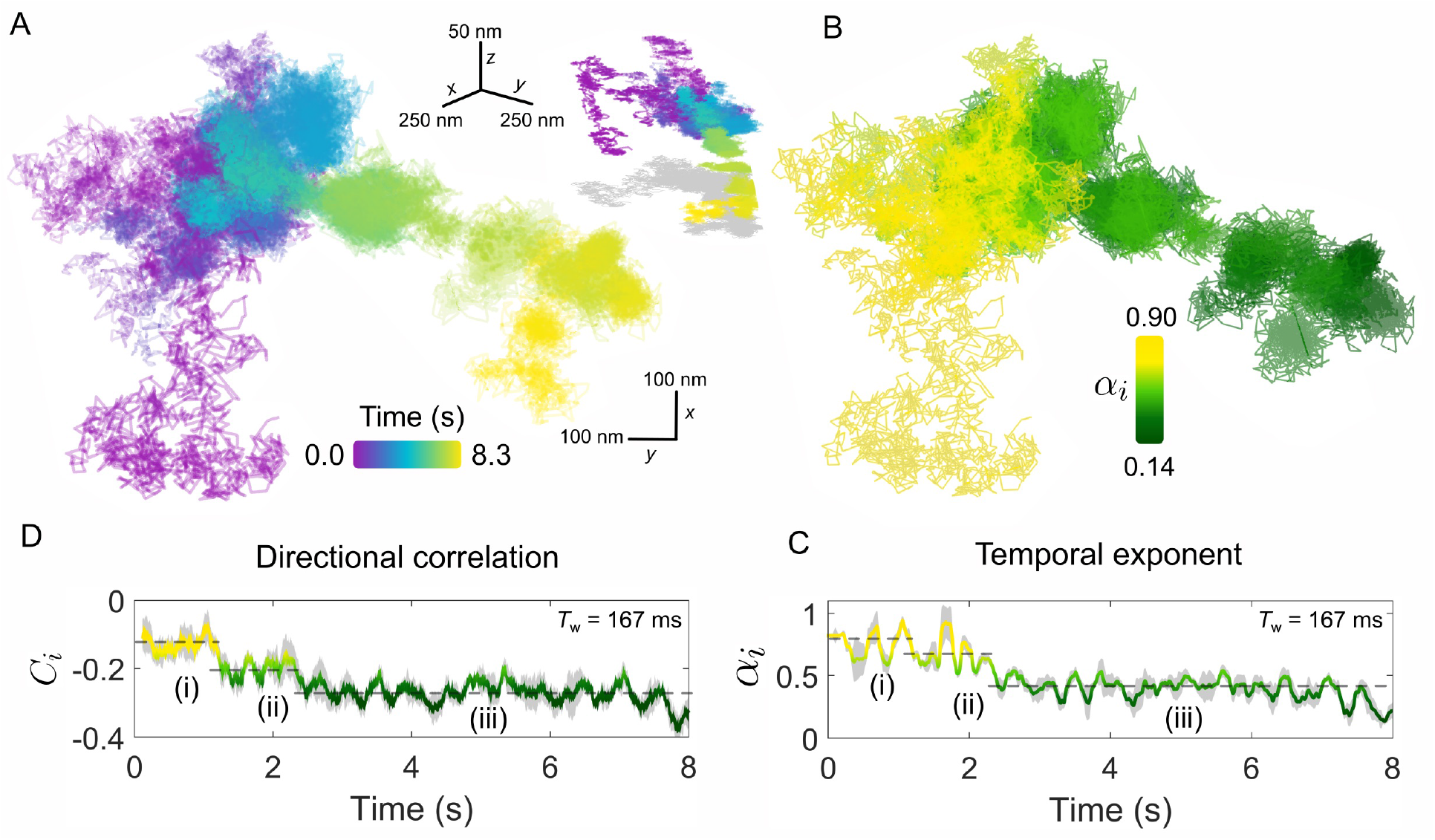
The endurance of iSPT imaging enables visualisation of dynamic, time-evolving processes. **(A)** An exemplar trajectory of a EGFR-GNP on a HeLa cell is plotted for a measurement over 8.3 s recorded at 6,000 fps. Inset: the 3D trajectory of the *x* -*y* projection shown in **(A)**, with a 2D ‘shadow’ projection plotted in grey. **(B)** The temporal exponent of the trajectory presented in **(A)** plotted spatially over the trajectory positions. **(C)** Temporal exponent plotted as an explicit function of time. **(D)** The corresponding directional correlation of the trajectory is also plotted as a function of time. Both temporal exponent and directional correlation use a sliding window of 1,000 frames (*T*_w_ = 167 ms) and the dashed lines guide the eye to the three distinct phases of diffusive behaviour, labelled (i)-(iii).

Such a change in the diffusional behavior of EGFR shown here is evocative of processes such as the binding of the EGFR-EGF complex to the actin skeleton beneath the membrane. The actin-binding domain located on the EGFR (78) can guide association with the cytoskeleton. This interaction has been shown to lead to an oriented transport by the flow of the skeleton accompanied by a decrease in diffusion (79), since remodelling of cortical actin is an active and important organisational motif for membrane molecules (80).

### Confined diffusion on the plasma membrane

Aside from sub-diffusion which still explores sufficiently large regions of the membrane, another mode observed is a diffusion where the probe moves very little from its original position and thus is often classified as being confined. These modes of diffusion are also interesting to investigate since restriction in the mobility of a membrane protein is an especially important aspect of membrane organisation and a means to regulate function of signalling proteins such as EGFR.

In **Fig. 8**, we present two examples of diffusion which is markedly confined within a spatial domain of approximately 100*×*100 nm^2^. The confined nature of the diffusion is most visually apparent when one compares the lateral trajectories of **Fig. 8(A,D)** with that of **Fig. 5(A)** which was recorded for the same duration and framerate. The temporal exponent of the diffusion, shown in **Fig. 8(B,E)** for a sliding window length of 1,000 frames (*T_w_* = 33 ms) similarly confirms the strong confinement of the EGFR with a consistent average value of *α* = 0.6 observed for both confined examples, although the trajectory of **Fig. 8(A)** shows slightly larger variance.

**Fig. 8.**
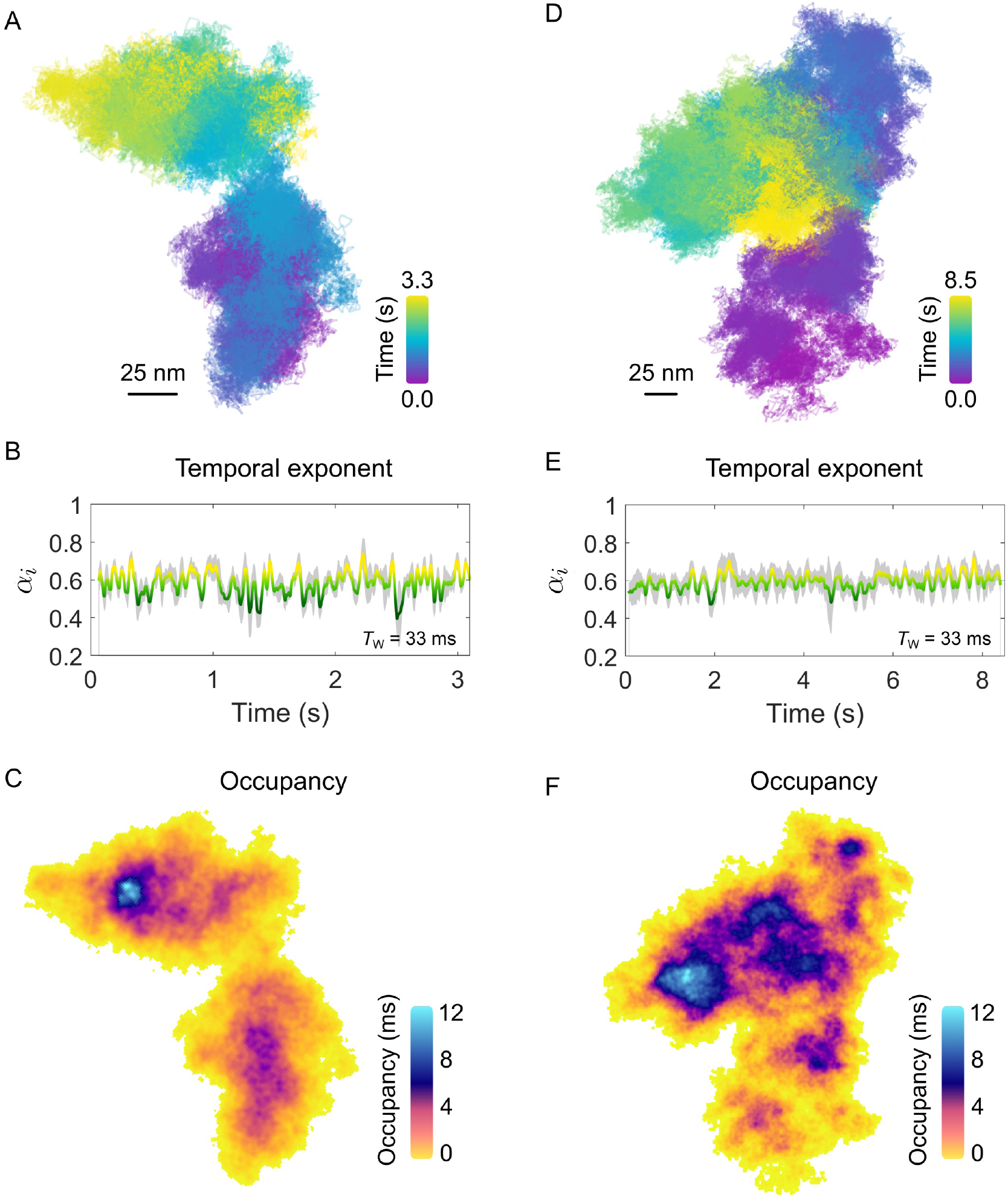
Confinement on the plasma membrane of a COS-7 cell. **(A)** The trajectory for an example of confined EGFR-GNP diffusion on a COS-7 cell is presented. The imaging framerate of 30,000 fps for a total of 3.3 s gives a total of 100,000 trajectory points. **(B)** The temporal exponent for the trajectory in **(A)** with an averaging window of *T*_w_ = 33 ms. **(C)** The respective ATOM plot for the trajectory of **(A)** is presented in **(C)**. The areal bin size is 4*×*4 nm^2^ and patches of extra-ordinary residence are reflected in the non-uniform colour scale encoding total occupation time. **(D)** The trajectory for a second example of confined membrane diffusion, recorded at the same framerate as **(A)**, but for 8.5 s, giving a total of 250,000 recordings. **(E)** The corresponding temporal exponent for the trajectory in **(D)** with an averaging window of *T*_w_ = 33 ms. **(F)** The ATOM plot for the trajectory of **(D)**, with an areal bin size is 4*×*4 nm^2^.

Here, in this example the diffusion occurs within a limited spatial region and within the apparent boundary of this region the entire space is repeatedly explored by the EGFR-GNP. This then opens the interesting possibility to examine the spatial occupancy of the trajectory through the ATOM plot wherein the frequent sampling of the membrane environment through our high-resolution tracking might reveal information about any structures influencing the EGFR-GNP diffusion. The ATOM plots for trajectories shown in **Fig. 8(A,D)** are plotted in **Fig. 8(C,F)** respectively for an areal 2D bin of 4*×*4 nm^2^. Inspection of the occupancy elucidated in the ATOM maps reveals distinct patches and heterogeneous patches of particularly long confinement. In **Fig. 8(B)** we identify a patch of *≈* 25 nm diameter as well as intermediate structures, typically located centrally, with also circular-like structures. In **Fig. 8(F)** we find numerous patches of extended occupation which are not homogeneous, but appear perforated with holes, suggesting regions of partial exclusion. Inspection also suggests the presence of circular-structures of a similar mean size of 25 nm, possibly pointing towards structures in the membrane the EGFR-GNP is interacting with whilst confined within the corral.

Another important example of confinement of a diffusing protein which occurs on the plasma membrane is that of confinement into ‘pits’ or ‘bowls’ that are often associated with endocytosis of membrane proteins. For example, it is know that one route for EGFR internalisation is through clathrin-mediated endocytosis, where clathrin assembles a bowl-like structure to envelope the portion of the membrane to be internalised.

In **Fig. 9(A,B)** we present an example of an EGFR-GNP trajectory, seen from two perspectives, recorded at 20,000 fps over 1.0 s, and a second example in **Fig. 9(C)** recorded at 30,000 fps over 3.2 s. Both examples show a 3D trajectory which reveals the EGFR-GNP is confined to the surface of a 3D bowl, with both examples presenting a bowl with an approximate diameter of 350 nm. The bowl morphology is most apparent when one interpolates through all the trajectory points to render an effective smooth surface, shown in **Fig. 9(D)**.

**Fig. 9.**
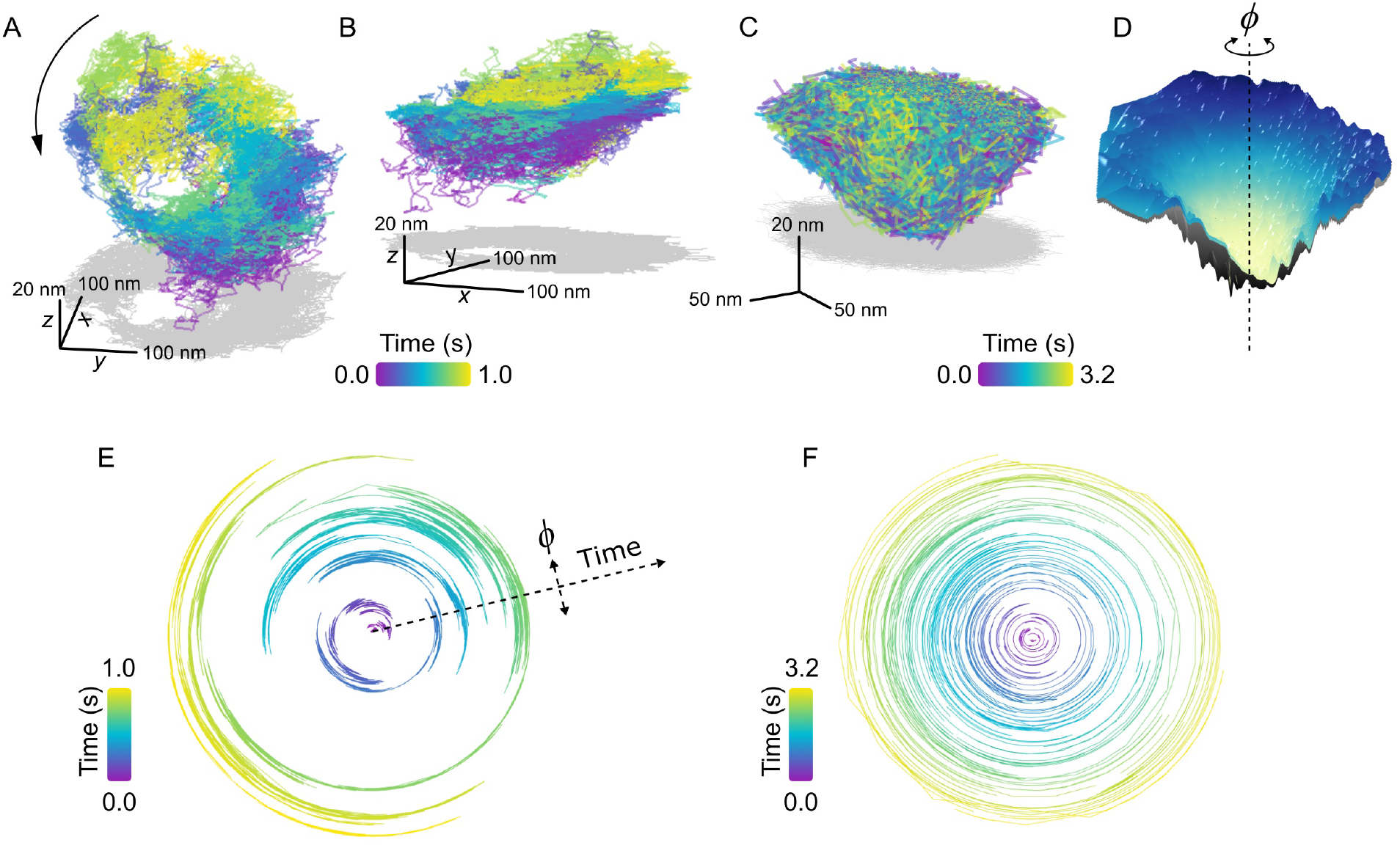
Confinement into bowl/pit-like on the plasma membrane. **(A,B)** Trajectory of a confined EGFR-GNP on a HeLa cell displaying a tilted ‘bowl’ or ‘pit’-like morphology, as seen from two perspectives. The trajectory was recorded at 20,000 fps for 1.0 s. Grey plots represents the 2D projection. **(C**) A second exemplary trajectory of a pit-like confinement, recorded at 30,000 fps for 3.2 s on a COS-7 cell. **(D)** An interpolated smooth surface of elucidating the pit topology from **(C**), with the centre of rotation marked with a dashed line. **(E)** Angular trajectory of the EGFR-GNP about the centre of the bowl from **(A)**, where the radial position denotes time. **(F)** Angular trajectory of the EGFR-GNP about the centre of the bowl from **(C)**.

In previous work, we observed the trajectory of the EGFR-GNP within such a bowl that suggested a persistent rotation of the probe around the centre of mass of the bowl. We denote the angular position of the probe with *φ*, shown schematically in **Fig. 9(D)**, and plot the angular position as a function of time for the trajectories of **Fig. 9(A)** and **Fig. 9(C)** in **Fig. 9(E)** and **Fig. 9(F)** respectively. In these angular plots, the radial extent denotes time, and in doing so one sees a back and forth rotation in **Fig. 9(E)**, as reported previously (30). It should be noted however, that not all bowl-like confinements necessarily exhibit this clear and persistent rotational motion. **Figure 9(F)** presents an example where no coherent rotational motion is seen, instead the probe was able to diffuse more erratically about the entire surface, but with occupancy favouring the lower regions of the bowl.

Given that EGFR is known to be internalised via clathrin mediated endocytosis and the similarity these EGFR-GNP bowl-trajectories bear with such pits, a potential biological origin is identified. It stands as an exciting line of future inquiry, however, to affirm the role these pits play in the EGFR membrane biology. Furthermore, one can identify whether the EGFR-GNP probe is mobile within a static membrane bowl, or whether the probe is bound to the membrane and the whole bowl itself undergoes rotation or alternatively whether a combination of the two is at play. Nonetheless, these measurements demonstrate the potential for iSPT to provide new avenues for deeper nanoscopic insight into established membrane biology.

#### Directed motion

A final example of another interesting mode of protein mobility is that of motion in a single concerted direction, as shown in **Fig. 10**. In the cellular context, such directed motion is typically due to active transport processes mediated by motor proteins that move organelles and endocytosed vesicles along the intracellular network of cytoskeletal filaments (81–83). We might suppose in the examples shown in **Fig. 10** that EGFR-GNP is internalised within a vesicle inside of the cell, and is being trafficked within the cellular corpus.

**Fig. 10.**
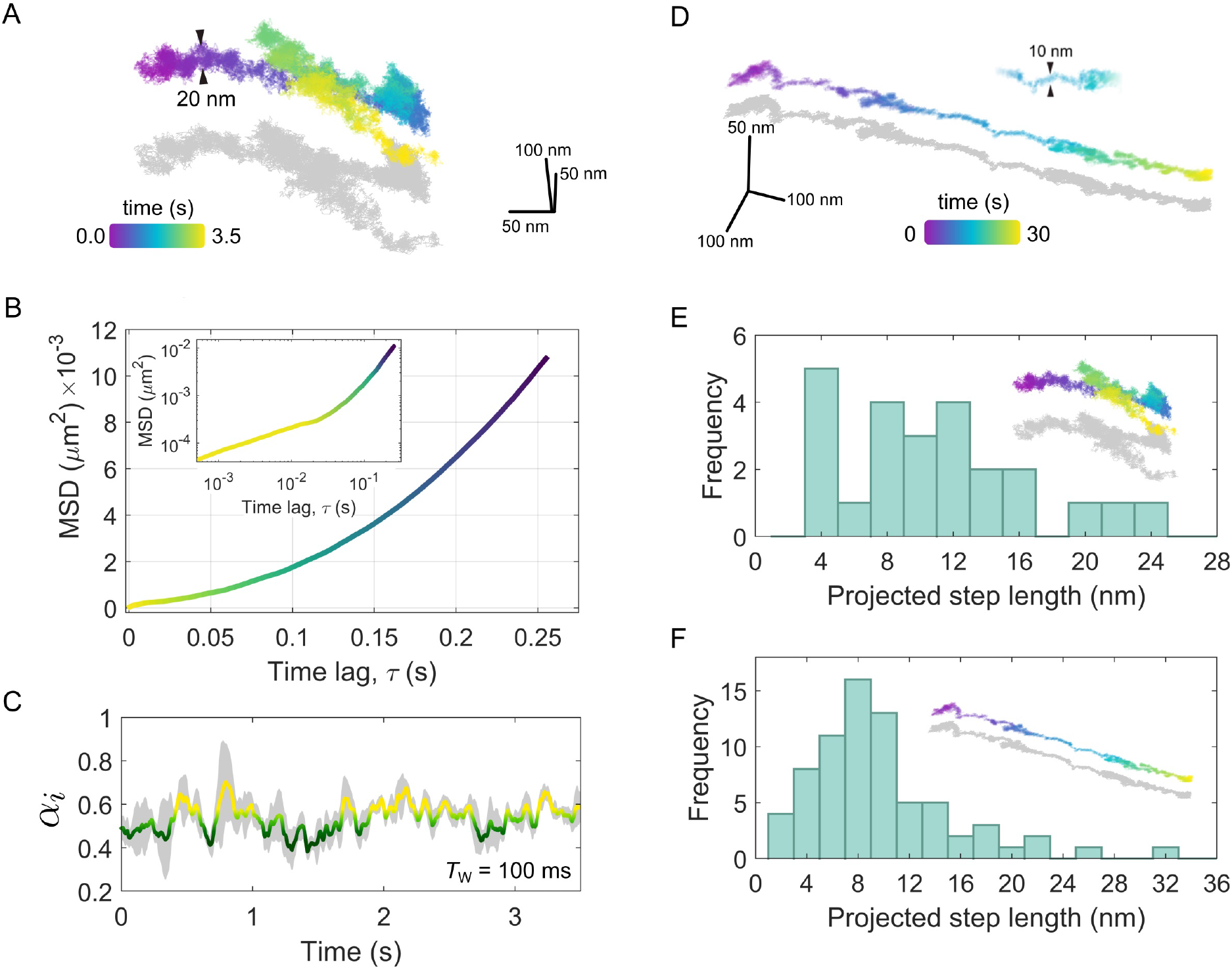
Directed motion of the EGFR-GNP probe. **(A)** A trajectory recorded at 10,000 fps for 3.5 s showing a concerted linear trafficking of the EGFR-GNP probe on a HeLa cell. The typical width of the trajectory track is 20 nm. **(B)** The MSD of the trajectory from **(A)**, showing super-diffusive mobility. The inset is the same MSD plotted on a log-log scale. **(C)** The temporal exponent of the trajectory from **(A)** for a rolling window of *T*_w_ = 100 ms. **(D)** A second example of a 3D trajectory showing directed motion, recorded at 1,000 fps for a duration of 30 s. The width of the track is approximately 10 nm. **(E)** A histogram of prominent steps projected into the direction of travel, from the trajectory in **(A)**. **(F)** A histogram of prominent steps projected onto the direction of travel, taken the trajectory shown in **(D)**.

In **Fig. 10(A)** we consider an example recorded at 10,000 fps wherein one sees a concerted diffusion along a given direction for several seconds that is then followed by a reverse walk and then a change in direction near orthogonal to the previous. Inspection of the trajectory in 3D reveals in fact that the second leg of the walk lies above the first, suggesting a switch in the track along which the walk occurs. The typical track width is seen to be around 20 nm.

A trajectory with such a persistent directionality is described as being super-diffusive as the time-dependence of the MSD as *α* > 1. In **Fig. 10(B)**, we present the macroscopic MSD computed by taking the complete trajectory at once, and find *α* = 1.98, which one can visualise through the upward curving slope of the MSD. The inset of **Fig. 10(B)** presents the same MSD plotted on a log-log scale wherein on can see clearly the super-diffusive transport behaviour manifest over longer time scales (*τ* > 0.01 *s*) and at shorter time scales (*τ* > 0.01 s) a more sub-diffusive time-dependence is apparent. We can explore this sub-diffusive aspect of the trajectory more closely by considering the rolling temporal exponent, shown in **Fig. 10(C)**, for a temporal window of *T_w_* = 100 ms. Here, one sees the temporal exponent fluctuates around *α ≈* 0.54. One can anticipate such ‘confined’-like behaviour in the short-time diffusion since the EGFR-GNP is likely confined to a vesicle which itself is tethered to the intracellular filament. Over longer timescales the processive mobility begins to emerge, reflected in the increasing value in *α*.

A second example, of super-diffusive mobility is presented in **Fig. 10(D)** for a trajectory recorded at 1,000 fps over 30 s observation time. In this example, a strongly linear track with a mean width of 10 nm is evident. Careful analysis of such trajectories shown in **Fig. 10(A)** and **Fig. 10(D)** allows us to not only identify the diffusive mode of transport but other important characteristics such as the velocity, step size and dwell time associated with the specific motor proteins which traffic cargo along these filaments within the cell. Taking into consideration that here we monitor the movement of EGFR most likely trapped in an intracellular vesicle, the inspection of these features of *in vivo* transport is fraught with complications and nuanced interpretation. In addition to active transport, velocity and step size will be affected by indirect movements such as polymerization or depolymerization of the filament it is attached to. Moreover, one must be mindful of the region within the cell where the EGFR-GNP is being tracked. If the filament is not lying flat to the coverslip, for example being located near the nucleus and orientated randomly, then what one observes in 2D is a projection of the true trajectory and hence it is not an accurate representation of the true step size. In iSCAT one can capture the 3D position so such a problem can be remedied. However, since the trajectories in **Fig. 10** appear near-parallel to the focal plane, one can assume this skew is not significant in these recordings. Whilst it is beyond the scope of this work to draw specific conclusions as to the behaviour exhibited in **Fig. 10**, it is nonetheless interesting to inspect what iSPT can provide in exploring *in vivo* transport and the issues surrounding it.

To inspect the occurrence of a preferred step length in the trajectories of **Fig. 10(A)** and **Fig. 10(D)** we define a linear axis that lies through the global direction of travel and onto this we project each individual step. We then may compute the accumulated distance walked by summing all projected steps and by using a peak-finding algorithm we can extract positions separating prominent steps along the walk. **Figure 10(E)** presents a histogram of the prominent projected steps from the trajectory shown in **Fig. 10(A)**(denoted by the inset) wherein one sees a preference for steps with an approximate integer multiple of 4 nm, most of which in a range up to 24 nm. In **Fig. 10(F)** a similar step-size histogram is given for the trajectory of **Fig. 10(D)**. In this latter example one also sees a similar distribution, but now with a stronger bias towards steps of *≈* 8 nm.

Elucidation of the step size associated with specific motor proteins has been the subject of numerous *in vitro* experiments wherein individual filaments and single motor proteins are investigated under pristine conditions e.g. (84–88). Whilst much of these investigations to quantify the step size have been done with fluorescence microscopies, recent efforts have harnessed the improved resolution of iSPT to identify the exact stepping mechanism of single myosin V (89, 90), also *in vitro*, which has remained a long standing matter of debate.

The tracking and interpretation of transport mediated by motor proteins *in vivo* within the living cell is a much more complex scenario, since it can involve more than just one transport mechanism. Various studies have, for example, revealed a switch of the transporting motor protein species such as from myosin, which travels on actin filaments to kinesin, which travels upon microtubules (91, 92). Additionally, these motor proteins not only alternate when transporting cargo, but can also be attached to the vesicle at the same time, requiring cooperative and even coordinated interactions (87, 93, 94). Furthermore, sub-division of the anticipated characteristic steps sizes can occur, as observed in *in vitro* experiments with kinesin (86) and in live cells with myosin II (95).

The step sizes and track widths of the trajectories in **Fig. 10** are in the range of step sizes reported for common molecular motor proteins including kinesin, myosin II, myosin V and VI in analogous *in vitro* and live cell measurements (86, 87, 95–98) and in good agreement with the widths of 7 nm of single actin filaments and 25 nm of microtubules (99). Therefore, these trajectories likely represent examples of active transport within the cell. However, lacking further information one may only speculate as to the molecular origin of these observed step sizes since the step size alone is not a distinctive characteristic of a particular motor protein and may be affected by interactions, cooperative and antagonistic effects in a living cell. One also has to consider that most of the *in vivo* and *in vitro* studies measure step size by labelling the motor protein itself, while we, in contrast, see the stepping behavior of the transported cargo.

To address the issue of intracellular transport more conclusively, one requires further investigation with a larger sample size to draw a more robust statistical analysis as well as measurement which harnesses fluorescence-based labelling to identify and visualise intracellular filaments and motor proteins in parallel with the iSPT tracking. Nonetheless, these initial efforts and recent similar *in vivo* iSPT tracking of cytoplasmatic vesicle transport (100) demonstrate that iSPT is well suited to investigate the nanoscopic minutia of intracellular transport events in living cells.

## Discussion: iSPT for investigation of mobility within the live cell

### iSPT microscopy

iSPT microscopy over the past few years has demonstrated great potential for new avenues of investigation into membrane diffusion, whether on synthetic or live cell membranes, owing to its high detection sensitivity affording fast and precise particle tracking. The power of the technique lies in the exploitation of interference between the light scattered by the probe and that which naturally reflects off the sample coverslip. For this reason iSPT, and more generally iSCAT microscopy, does not require any special equipment, save for a suitably fast camera to achieve high imaging rates. This makes iSCAT microscopy easy to implement on existing microscopes, including commercial fluorescence microscopes, and as both microscopies are mutually compatible (56) they can also be performed in parallel. This is particularly important in live cell imaging where fluorescent labelling is useful for identifying specific cell and membrane components and features.

### Phototoxicity

One important issue in live cell imaging is the role of photodamage associated with high laser intensities commonly associated with high-resolution microscopies (101). The threshold for potential cell damage depends on the particular cells in use as well as the particular aspect of the membrane and cell biology under investigation (55). We have demonstrated previously one can achieve high framerate and accurate tracking of EGFR-GNP probes on HeLa cells using illumination powers that are tolerable for cell viability (30), being on the order of no more than 10 kWcm^*−*2^. In general, using illumination wavelengths towards the red end of the spectrum (beginning around 600 nm) appears to be the most compatible for preserving cell viability. One asset of iSCAT microscopy is that, unlike fluorescence, one has complete flexibility in choice of the wavelength used and thus it is possible to perform iSCAT microscopies at wavelengths that preserve cell health whilst also maintaining high framerate imaging and good signal-to-noise ratio.

### Influence of the probe

An issue often raised in SPT is to what extent the probe influences the diffusive process and membrane biology under investigation. The label, depending on its geometry and functionalisation introduces multivalent binding, non-specific binding as well as the physical size adding steric hindrances to the crowded membrane environment. The latter of which is an important consideration for colloidal probes particularly used for iSPT. In addition, because the localisation precision now can approach the few-nm length scale, the influence of the GNP-to-ligand linker becomes of interest, in particular its length and flexibilty, and how these might affect the accuracy in localisation of the associated protein. Many of these questions are not fully answered, but recent studies have suggested that the size of these colloidal probes may not have significant effects on diffusion within the cell as one would have initially assumed (54, 102, 103).

Similarly, in previous work, we also found the use of 48 nm EGF-GNP probes seemingly did not impair the EGFR signalling pathway or uptake via clathrin-mediated endocytosis (30). To fully address this matter, one must systematically explore membrane diffusion for colloidal probes of differing sizes, as it has been recently demonstrated for synthetic membranes (104). Usage of smaller colloidal probes on the live cell membrane is challenging solely due to the dynamic speckle background which presents the biggest hurdle to overcome. However, new methods for background modelling and subtraction in interferometric microscopies are constantly being proposed (30, 35, 60, 105), and so it is realistic that the challenge posed by the cell background will be met within the near future.

## Conclusion and Outlook

iSPT microscopy stands as a promising technique to expand the frontiers of investigation into single molecule diffusion and membrane organisation. The high resolution visualisation of protein mobility in 3D provides a wealth of information hitherto inaccessible through conventional fluorescence microscopies. When coupled with complementary techniques such as super-resolution fluorescence and correlative electron microscopies, one is presented with an impressive tool box that could help addressing long standing questions regarding dynamic membrane organisation such as the size, lifetime and diffusive properties of membrane raft domains.

Furthermore, more broadly, iSCAT microscopy is an emergent and rapidly growing technique that is finding increasing applications in nanoscale biology, with aims ranging from label-free detection and mass-determination of single proteins and complexes, to the detection of viruses and vesicles as well as live cell imaging. Recent interdisciplinary efforts to model the iPSF as well as to address dynamic speckle background through machine learning opens new paths to exciting live-cell applications. This will allow the use of smaller scattering probes, multiple distinct scattering labels for co-labelling investigations as well as being able to perform imaging and iSPT over extended axial ranges within deeply scattering tissues.

## Materials and Methods

### Widefield iSCAT microscopy

Laser light from a supercontinuum white-light laser (NKT Photonics) was filtered down to *λ*_iSCAT_ = 550 *±* 15 nm through a Varia filter box. An 100x oil-immersion (NA = 1.4) objective was used to give a field of view of 5 *×* 5*μ*m^2^, which was then imaged onto 128×128 pixels of a high-speed camera (Vision Research, Phantom, Miro LAB 3a10).

In practice, illumination can be from any coherent light source, for gold nanoparticles green light is optimal. The illumination is unpolarised to minimise polarisation artefacts which can arise for high-NA imaging. The objective is oil-immersion to maximise collection efficiency.

### Confocal fluorescence imaging

A Zeiss LSM 800 was used with a water-dipping objective (40x), and modified to accommodate an iSCAT microscope underneath. A wavelength of 650 nm was chosen to avoid overlapping with the wide-field iSCAT imaging wavelength and to mitigate potential damage to live cells.

### Cell culturing & fluorescent labelling

HeLa and COS-7 cells were grown in DMEM (Gibco Invitrogen) supplemented with 10% fetal calf serum (FCS, Life Technologies) in a humidified atmosphere at 37°C and 5% CO_2_. For measurement, HeLa cells were plated onto a glass-bottomed sample dish (Ibidi GmBH) and grown to 70% confluency. Before measurement, each dish was rinsed twice with warmed PBS-BSA solution, serum starved for several hours and rinsed again with warmed PBS-BSA. Imaging was done in Leibovitz’s L-15 Medium (1.5 ml Gibco, Invitrogen).

For tracking over-expressed EGFR on COS-7 cells, cells were counted and 90,000 cells per dish were seeded. The next day, medium was replaced by 3 ml fresh DMEM and the Lipofectamine 3000 transfection mix was added (prepared according to the manufacturer’s protocol (ThermoFisher). (Details:4.6 *μ*l Lipofectamine 3000, 2 *μ*g plasmid DNA (EGFR-EGFP, addgene #32751) and 4 *μ*l P3000 Reagent). For imaging, cells were rinsed twice with DPBS and 3 ml of pre-warmed Leibovitz’s L-15 medium (without phenol red, ThermoFisher) were added.

When placed on the microscope, each culture was held at 37°C by a home-built objective heater. A micropipette (Piezo Drill Tip, Eppendorf) was used to deliver 10 *μ*l of the EGF–GNPs to a local region of the culture, with observation beginning immediately thereafter.

### Gold nanoparticle probes

GNPs with a diameter of 48 nm were functionalised with monoclonal anti-biotin (British Biocell International), and GNPs with diameters of 20 nm, functionalised with streptavidin (British Biocell International), were conjugated at a molar ratio of 1:1 with biotinylated EGF (ThermoFisher) at a concentration of 0.66 nM. PBS was used as a buffer. The solution containing EGF–GNP was agitated for several hours at 37°C to assist conjugation, purified through centrifugation and diluted up to to a final concentration of 0.1 nM. Cationic GNPs with diameter 50 nm were purchased from Nanopartz Inc. (item CC11-50-POS-DIH-50-1) and diluted to a ratio 1:100 in water.

## Author Contributions

R.W.T. and C.H. made iSCAT and fluorescence measurements. R.W.T. performed data analysis and C.H. prepared the live cell materials. R.G.M. and H.M.D. developed background correction routines for the iSCAT images. M.K. developed the combination of confocal fluorescence and iSCAT microscopy. V.Z. supervised interpretation and analysis of the data. A.S. supervised biological preparation and data interpretation. V.S. conceived and supervised the project. R.W.T., C.H., V.Z. and A.S. wrote the manuscript. All authors discussed the results and commented on the manuscript.

## Acknowledgments

This project was funded by an Alexander von Humboldt professorship and postdoctoral fellowship as well continuous support from the Max Planck Society. The authors wish to thank Jennifer Lühr for assisting with the development of GNP pipetting, Moritz Grob and David Albrecht for careful reading of the manuscript and insightful comments, Simone Ihloff for assistance with the culturing of cells, Navid Bonakdar, Maksim Schwab, Oliver Bittel and Lothar Meier for designing and fabricating the sample heating system.

